# A solution to the biodiversity paradox by logical deterministic cellular automata

**DOI:** 10.1101/006817

**Authors:** Lev V. Kalmykov, Vyacheslav L. Kalmykov

## Abstract

The paradox of biological diversity is the key problem of theoretical ecology. The paradox consists in the contradiction between the competitive exclusion principle and the observed biodiversity. This principle was formulated incorrectly because of limitations of the traditional black-box models of interspecific competition. The principle is very important as the basis for understanding evolutionary processes. Our white-box multiscale models are based on logical deterministic individual-based cellular automata. This approach allows to provide an automatic deductive inference on the basis of a system of axioms and to get a direct holistic insight into the studied system. It is one of the most promising methods of artificial intelligence. Here on simplest models we show a mechanism of competitive coexistence which violates the known formulations of the competitive exclusion principle. We reformulate and generalize the competitive exclusion principle and explain why our formulations provide a solution of the biodiversity paradox. In addition, we propose a principle of competitive coexistence.

**Graphical abstract:** 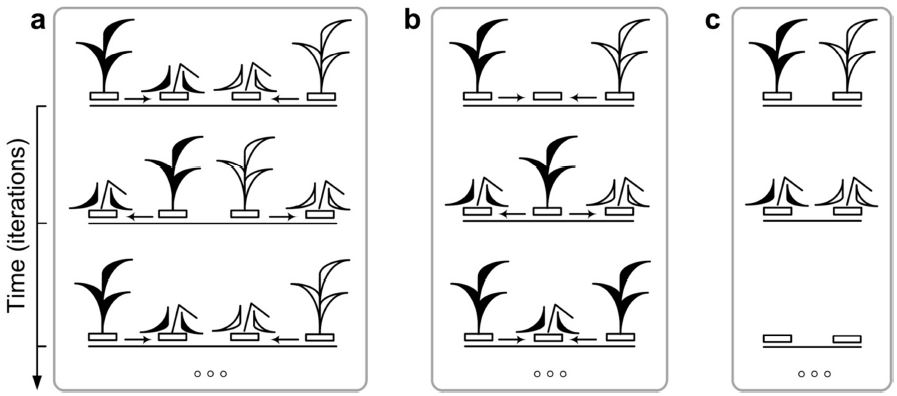

## 1. Introduction

There is a long-standing question in theoretical ecology – How so many superficially similar species coexist together? (Craze, 2012; Palmer, 1994; Sommer, 1999) Trying to answer this question, ecologists came to the biodiversity paradox (Hutchinson, 1961; Lehman and Tilman, 1997; Sommer, 1999). In theory, according the competitive exclusion principle, complete competitors cannot coexist (Hardin, 1960). In practice there are many examples of such coexistence: rain forests (Hubbell, 2001), coral reefs (Dornelas et al., 2006), grasslands (Tilman et al., 2006), plankton communities (Hutchinson, 1961; Sommer, 1999). The paradox consists in the contradiction between the competitive exclusion principle and the observed natural biodiversity. “The apparent contradiction between competitive exclusion and the species richness found in nature has been a long-standing enigma” (Sommer, 1999). “Resolving the diversity paradox became the central issue in theoretical ecology” (Lehman and Tilman, 1997). The key question in this long-standing problem is a validity of the competitive exclusion principle. Is this principle true? “There are many who have supposed that the principle is one that can be proved or disproved by empirical facts among them Gause himself. Nothing could be farther from the truth. The “truth” of the principle is and can be established only by theory, not being subject to proof or disproof by facts, as ordinarily understood” (Hardin, 1960). Competitive exclusion principle is very important as the basis for understanding evolutionary processes. It postulates the mechanism by which a more adapted species wins in the struggle for existence.

We assume that the difficulties of verifying the competitive exclusion principle are associated with limitations of the traditional black-box models of complex systems. We consider a complex system as consisting of subsystems, which are hierarchically subdivided into micro-, meso-subsystems and a whole macro-system. Interactions between subsystems of a complex system lead to emergence of new properties on the macro-level that are absent on the micro- and meso-levels. In our opinion, logical deterministic cellular automata are the most appropriate tool for investigation of a complex system in time and space. Such approach is bottom-up mechanistic and provides a direct visual insight into dynamics of the studied complex systems on micro-, meso- and macro-levels. It provides a strong white-box and really multiscale modelling of complex systems. Only a model that is based both on cause-and-effect and part-whole relationships between subsystems of a complex system may help to achieve a holistic understanding of how this system works. Our study focuses on modelling of population and ecosystem dynamics. Earlier we create white-box models based on logical deterministic individual-based cellular automata that simultaneously consider both part-whole and cause-effect relationships (Kalmykov and Kalmykov, 2013). Here we give an explicit specification of this approach and use it to solve the biodiversity paradox in models of smallest ecosystems.

## 2. Hypothesis: complete competitors can coexist

We suppose that complete competitors which are identical consumers and differ only in fitness can indefinitely coexist in one very small closed homogeneous habitat on one and the same limiting resource under constant conditions of environment, without any trade-offs and cooperations. A mechanism of such coexistence may be based on timely recovery of essential resource.

## 3. Theory and method

In order to test this hypothesis we have created and investigated simplest, visual and self-evident mathematical models of population and ecosystem dynamics. The simplest models were used because the competitive exclusion principle must be equally valid both for complex and for simple cases. The simplest cases are more preferable as they are the most evident. These models are logical deterministic individual-based (i.e. multi-agent) cellular automata. Pure logical modelling with detailed visual representation of all stages of ecosystem dynamics allows us to interpret results strictly mechanistically. All states of a lattice site and transitions between these states have mechanistic interpretations. It provides correct interpretations of the results. Discrete time, discrete space and deterministic rules make such models clear for mechanistic understanding.

Deterministic logical individual-based cellular automata models of complex systems are of white-box type. A white box model has “transparent walls” and directly shows underlined mechanistic mechanisms – all events at micro-, meso- and macro-levels of the simulated dynamic system are directly visible at all stages (Kalmykov and Kalmykov, 2013). The first step of the white-box modelling of a complex system is a top-down creation of a set of axioms on the basis of holistic understanding of a domain under study. The second step is creation of a white-box model of the domain by automatic logical bottom-up inference from the formulated set of axioms. These two steps reflect the top-down-bottom-up nature of the white-box modelling of a complex system.

There are three types of mathematical models of complex systems: black-box, grey-box and white-box models. Figurative representation of these models is presented in Fig.1. Differential equation, stochastic and matrix models do not model individuals and their local interactions. That is why we consider them as non-mechanistic black-box models which cannot provide a direct insight into a complex system dynamics.

**Figure 1.**
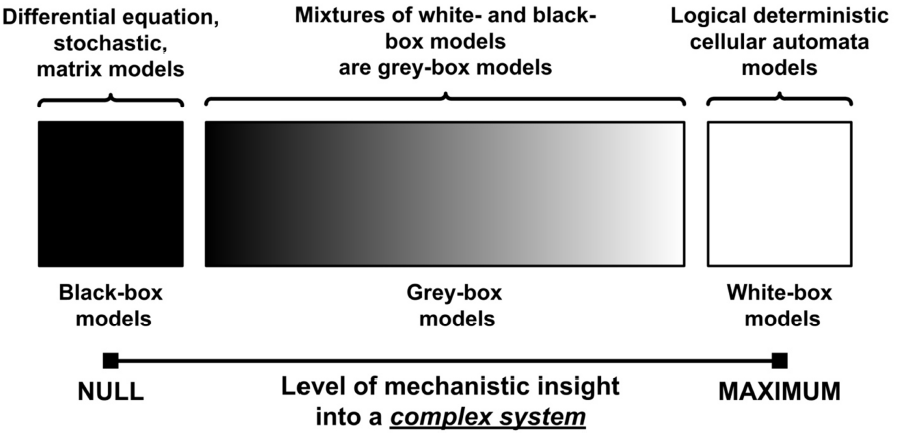
Three types of mathematical models of complex systems. For example, Verhulst model of population growth, Lotka-Volterra model of interspecific competition and Lanchester models of warfare are black-box models. These models are phenomenological and non-mechanistic (Tilman, 1987). Stochastic cellular automata are examples of grey-box models. Deterministic logical axiomatic cellular automata models of complex systems are of white-box type (Kalmykov and Kalmykov, 2013).

The key stage of the cellular automata modelling of a complex system is a formulation of physically adequate axioms. These axioms must represent the highest possible degree of physical generalization of a studied complex system. The necessity of adequate basic understanding of a research area is the main difficulty here. A system of axioms created on the basis of this fundamental understanding is used for formulating rules of automatic logical deterministic inference of white-box models. Here we introduce a physics-based semantics (ontology) of our ecosystem models. Further, we transform this semantics to a set of axiomatic cellular automata rules. An elementary object of our models is a microecosystem (Fig. 2). Following Tansley we consider individual with its “special environment, with which they form one physical system” (Tansley, 1935).

**Figure 2.**
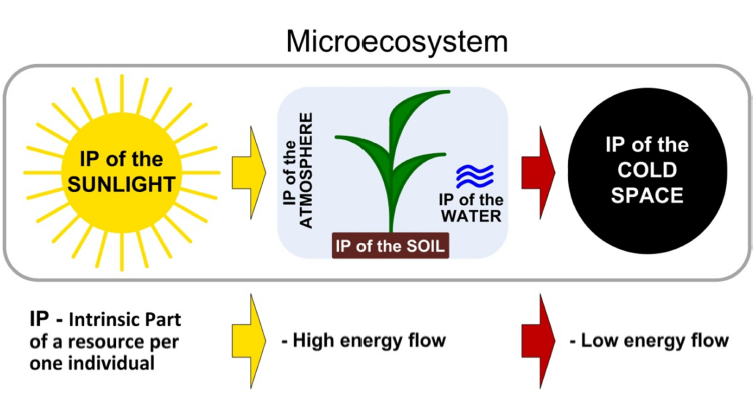
Schematic representation of a microecosystem as a microhabitat with an individual living in it. A microhabitat is the intrinsic part of environmental resources of one individual and it contains all necessary resources for its life. The high-grade (“fresh”) energy flow of solar radiation provides an individual’s life in a microhabitat, and the flow of lowered-grade (“used-up”) energy which produced during an individual’s life goes into the cold space.

A microhabitat includes the totality of resources and conditions that are necessary for a single individual’s life. It is intrinsic part of all environmental resources and conditions per one individual. A living organism is not able to function without the heat output to a “refrigerator”. Ultimately, the cold space performs a role of the such refrigerator. A working cycle of microecosystem’s processes represents a closed circle, which includes occupation of a microhabitat by an individual and restoration of used resources after an individual’s death (Fig. 3d,e). An ecosystem consisting of a single microhabitat is able to provide a single accommodation of one individual but it cannot provide its propagation. This is due to the fact that after an individual’s death, its microhabitat goes into regeneration (refractory) state, which is not suitable for immediate occupation. A minimal ecosystem, providing a possibility of vegetative propagation of an individual must consist of two microhabitats. In our models vegetative propagation is carried out due to resources of an individual’s microhabitat by placing finished vegetative primordium of descendant in a free microhabitat of an individual’s neigbourhood. A feature of our white-box modelling of ecosystems dynamics is the closure of all interactions of an individual with its intrinsic micro-environment at the level of microecosystem. This closure allows to consider a microecosystem as autonomous object. Spontaneous dynamics of microecosystem modelled as transitions between the states of a site. These transitions are carried out in accordance with extreme principles of physics.

Here we describe two of our white-box models. The first model represents an ecosystem with one species. The second one represents an ecosystem with two competing species.

Mathematically these models are cellular automata and can be defined by the group of five interrelated objects:

1. A cellular automata lattice, uniting a collection of sites;
2. A finite set of possible states of a lattice site;
3. A cellular automata neighbourhood which consists of a site and its intrinsically defined neighbour sites;
4. A function of transitions between the states of a lattice site;
5. An initial pattern of the states of all lattice sites.

An each lattice site is a finite automaton. Changing of the states of each site is determined according to the transition function. States of each of all sites are iteratively updating synchronously at discrete time steps. The transition function determines the state of each lattice site, at the next iteration according to logical rules. An iteration of the cellular automata consists of application of the transition function to each of all lattice sites. The neighbourhood of a site is a specific structure and it is defined as the site itself and a special set of its neighbouring sites.

Basic provisions of the models:

1. The whole cellular automata model simulates an ecosystem;
2. A microecosystem modeled by a lattice site;
3. A possible spatial pattern of vegetative propagation and the maximum number of offsprings per one individual are determined by the neighbourhood;
4. A free microhabitat contains resources and conditions for an individual’s life;
5. An occupied microhabitat goes into the regeneration state after an individual’s death;
6. The used resources of a microhabitat are recovered after living of an individual during one next iteration (i.e. during the regeneration state of a site);
7. A microhabitat may be occupied immediately after finishing of the regeneration state.
8. A populated microhabitat and a microhabitat in the regeneration state can not be occupied;
9. A neighbourhood determines a possible spatial pattern of vegetative propagation and the maximum number of offsprings per one individual;
10. Individuals are immobile in the lattice sites;
11. The lattice is closed. It may be closed on the torus by periodic conditions or by boundary conditions on the plane.

### 3.1. White-box models of population and ecosystem dynamics

Here we characterise our cellular automata models of population and ecosystem dynamics in more details. We start from an ecosystem model with one species and then investigate the ecosystem model with two competing species. We present an ecosystem model with a one species in Fig. 3. A two-dimensional lattice is closed on the torus (Fig. 3f,g). The neighbourhood can be of any type, but here we use only the hexagonal one (Figs. 3a and 4a). The neighbourhood allows to model a vegetative propagation of plants when an individual’s offsprings can occupy all nearest microhabitats (Fig. 3b, c). The hexagonal neigbourhood provides the closest packing of individuals.

Here is a description of states of a site in the ecosystem model with a single species. Each site may be in one of the three states 0, 1 or 2, where:

0 – a free microhabitat which can be occupied by an individual of the species;
1 – a microhabitat is occupied by a living individual of the species;
2 – a regeneration state of a microhabitat after death of an individual of the species.

A transition function includes logical rules of transitions between the states of a site of the single-species model. Let’s list these logical rules (Fig. 3d,e):

0→0, a microhabitat remains free if there is no one living individual in its neighbourhood;
0→1, a microhabitat will be occupied by an individual of the species if there is at least one individual in its neighbourhood;
1→2, after death of an individual of the species its microhabitat goes into the regeneration state;
2→0, after the regeneration state a microhabitat becomes free if there is no one living individual in its neighbourhood;
2→1, after the regeneration state a microhabitat is occupied by an individual of the species if there is at least one individual in its neighbourhood.

A set of possible states of a lattice site and a transition function can be represented by the directed graph of transitions between the states a lattice site (Fig. 3d,e). A pictorial representation (Fig. 3b,c,e,f) is given to facilitate understanding. An example of initial pattern is represented in Fig. 3f,g in two forms. There is a single individual on the lattice consisting of 5×5 sites. Fig. 3f,g shows spatio-temporal patterns of ecosystem’s colonisation by one species. Fig. 3h shows population dynamics. Increasing of the lattice size results in a logistic-like S-curve (Fig. 3i). Here we mechanistically investigated the formation mechanism of this curve. This mechanistic insight cannot be achieved by using black-box models. All steps of population and ecosystem dynamics are visualized up to the destiny of each separate individual.

**Figure 3.**
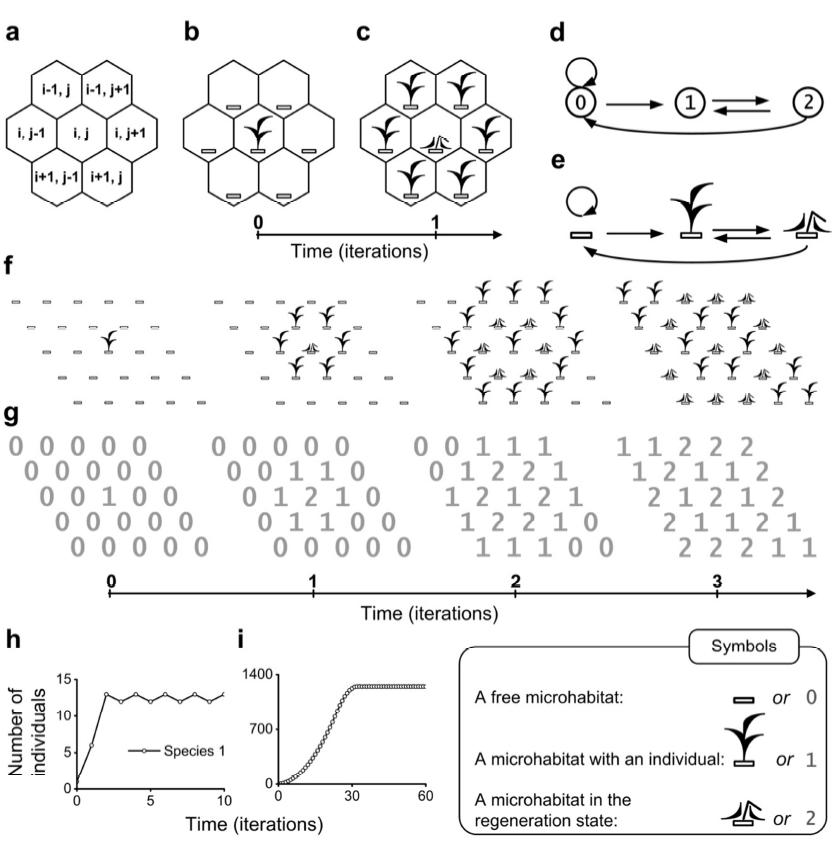
An ecosystem model with a single species. **a**, Hexagonal neighbourhood where i and j are integer numbers. **b,c,** Propagation of an individual in the hexagonal neighbourhood. **d,e**, Directed graph of transitions between the states of a lattice site of the ecosystem model with a single species. This graph has a numerical (d) and pictorial (e) representations. **f**,**g,** Spatio-temporal patterns of the model are represented in two forms: pictorial (f) and numerical (g). The lattice consists of 5×5 sites. Individuals propagate in the hexagonal neighbourhood. **h**, Population dynamics of the model (f,g) during 10 iterations. **i**, Population dynamics during 60 iterations. The lattice consists of 50×50 sites.

We implemented the accordance of our ecological formalism with the classic axiomatic formalism of Wiener and Rosenblueth for simulation of excitation propagation in active media. Three successive states – rest, excitation and refractoriness of each site are the main features of that formalism. In our formalism the ‘rest’ state corresponds to the ‘free’ state of a microhabitat, the ‘excitation’ corresponds to the life activity of an individual in a microhabitat and the ‘refractoriness’ corresponds to the regeneration state of a microhabitat (Fig. 3d,e). It simulates a birth–death–regeneration process in an ecosystem. This process implements the ecosystem’s working cycle as the working cycle of a “heat engine” mechanism, in which the “heater” is the sun, and “refrigerator” is the cold space (Fig. 2).

### 3.2. A model of interspecific competition

Here we describe our cellular automata model of interspecific competition. A finite set of possible states of the lattice site 0, 1, 2, 3, 4, 5 (Fig. 4b). A neighbourhood is hexagonal (Fig. 4a). A function of transitions between the states of a lattice site is represented by directed graph (Fig. 4b,c). The transition graph is represented in two forms: numerical and pictorial. Initial patterns of the states of a lattice site are defined at initial iteration of the model (Fig. 5).

**Figure 4.**
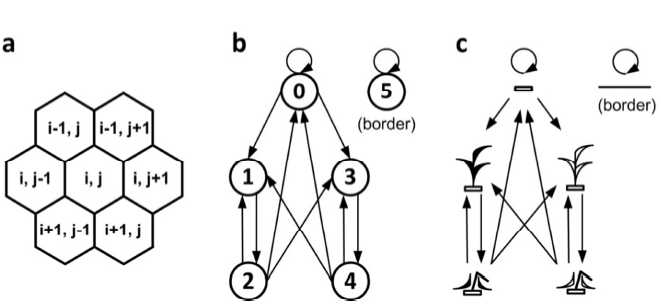
An ecosystem model of two-species competition. **a,** Hexagonal neighbourhood where i and j are integer numbers. **b-c**, Directed graph of transitions between the states of a lattice site. The graph has a numerical (b) and pictorial (c) representations. Individuals of the dominant species 1 are black (c). Individuals of the recessive species 2 are white (c). Individuals of the competing species differ only in fitness.

Description of the states of the two-species competition model (Fig. 4b,c). Each site may be in one of the five states:

0 – a free microhabitat which can be occupied by an individual of the both species;
1 – a microhabitat is occupied by a living individual of the first species;
2 – a regeneration state of a microhabitat after death of an individual of the first species;
3 – a microhabitat is occupied by a living individual of the second species;
4 – a regeneration state of a microhabitat after death of an individual of the second species;
5 – a site in this state represents an element of the boundary conditions, i.e. border.

Logical rules of transitions between the states of a site of the two-species competition model (Fig. 4b,c):

0→0, a microhabitat remains free if there is no one living individual in the neighbourhood;
0→1, a microhabitat will be occupied by an individual of the first species if there is at least one individual of the first species in its neighbourhood;
0→3, a microhabitat will be occupied by an individual of the second species if there is at least one individual of the second species in its neighbourhood and there is no one individual of the first species in its neighbourhood;
1→2, after death of an individual of the first species its microhabitat goes into the regeneration state;
2→0, after the regeneration state a microhabitat will be free if there is no one living individual in the neighbourhood;
2→1, after the regeneration state a microhabitat will be occupied by an individual of the first species if there is at least one individual of the first species in its neighbourhood;
2→3, after the regeneration state a microhabitat will be occupied by an individual of the second species if there is at least one individual of the second species in its neighbourhood and there is no one individual of the first species in its neighbourhood;
3→4, after death of an individual of the second species its microhabitat goes into the regeneration state;
4→0, after the regeneration state a microhabitat will be free if there is no one living individual in its neighbourhood;
4→1, after the regeneration state a microhabitat will be occupied by an individual of the first species if there is at least one individual of the first species in its neighbourhood;
4→3, after the regeneration state a microhabitat will be occupied by an individual of the second species if there is at least one individual of the second species in its neighbourhood and there is no one individual of the first species in its neighbourhood;
5→5, a site remains in this state, which means the border of the ecosystem.

The ecosystem is closed on the plane by the boundary conditions. The border modelled by the specific state of a lattice site. The state ‘5’ of a site denotes the border. Modelling of ecosystem dynamics on the plane is more natural than on the torus. Introduction of this boundary condition provides an extension of our earlier proposed model of ecosystem with interspecific competition between two species (Kalmykov and Kalmykov, 2013).

## 4. Results and discussion

Here, we automatically verify our hypothesis using the model of interspecific competition. We consider very small ecosystems which consist of two, three and four microhabitats with single individuals of the two competing species (Fig. 5). This approach simultaneously and holistically combines pictorial representation and mechanicalness by uniting determinism, logic and discreteness of time and space. On the basis of the logical rules we investigated patterns with different initial conditions, such as the lattice size and initial positioning of individuals on the lattice. We investigated four key cases of ecosystem dynamics and present results in two versions. The first presentation is pictorial (Fig. 5a–d) and the second one represents the same cases but in numerical form of program implementation (Fig. 5e–h). We model two complete competitors, which differ only in fitness. Fitness is defined here as a primary ability of an individual of a species to occupy a free microhabitat in a direct conflict of interest with an individual of another species (Figs. 5a,e). A dominant species wins in the direct conflict of interest in 100% of cases. These model conditions exclude any interspecific trade-offs and cooperations. In Fig. 5a,e we show a simplest case of competitive exclusion in result of the direct conflict of interest between individuals of the competing species. Now, we pass to the verification of our hypothesis. In Fig. 5b,f we prove that complete competitors which differ only in fitness can indefinitely coexist in one closed homogeneous habitat on one and the same limiting resource under constant conditions of environment, without any trade-offs and cooperations. A mechanism of such coexistence is based on timely recovery of essential resource, structural features of the habitat and initial positioning of individuals of the competing species. This is a clear and simplest proof of violation of the earlier formulations of the competitive exclusion principle which postulated that “complete competitors cannot coexist” (Hardin, 1960). There is no doubt that the principle should be equally valid for all ecosystems - including the smallest. Our previous results also revealed a violation of the competitive exclusion principle and we were forced to reformulate this principle as follows (Kalmykov and Kalmykov, 2013):

> If each and every individual of a less fit species in any attempt to use any limiting resource always has a direct conflict of interest with an individual of a most fittest species and always loses, then, all other things being equal for all individuals of the competing species, these species cannot coexist indefinitely and the less fit species will be excluded from the habitat in the long run.
>
> — *(Definition 1)*

*Interspecific resource competition* is a competition between individuals of different species for the same limiting resources in one ecosystem. Under the *limiting resource* we mean a *scarce essential resource*, which directly limits a reproduction of individuals of the species. The *essential resource* is a resource that is fundamentally necessary for existence, functioning and reproduction of individuals of the species and that cannot be replaced by another resource of the species’s intrinsic ecosystem. Fig. 5a,e demonstrates the case of implementation of the competitive exclusion principle (Def. 1) in result of the direct conflict of interest between individuals of the competing species. The species 1 wins the species 2. This is the smallest ecosystem for realization of competitive exclusion in result of the direct conflict of interest. It consists of the three microhabitats and two individuals of the competing species.

**Figure 5.**
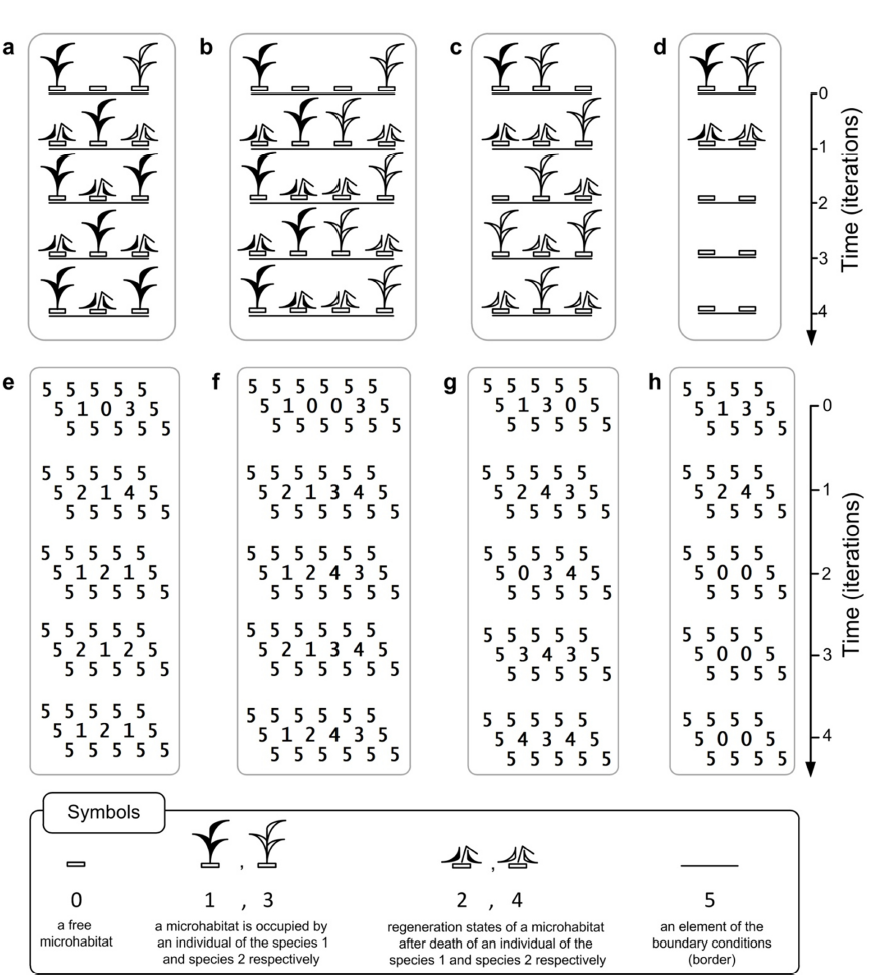
Verification of the hypothesis that “complete competitors can coexist”. Spatio-temporal patterns of the two-species competition model are represented in two forms: pictorial (a–d) and numerical (e–h). The species 1 has greater fitness than the species 2 and this is the only one difference between them. **a**,**e**, The case of competitive exclusion in result of the direct conflict of interest between complete competitors. **b**,**f**, Complete competitors indefinitely coexist in one closed homogeneous habitat on one and the same limiting resource under constant conditions of environment, without any trade-offs and cooperations. **c**,**g**, Paradoxical case of competitive exclusion. The species 2 wins the species 1 in result of indirect competition. **d**,**h**, The both competitors extinct due to the lack of available resource. The both competitors cannot propagate.

According to the Definition 1 an implementation of the competitive exclusion principle in nature requires too strict conditions. Implementation of the competitive exclusion of a species is especially improbable in tropical rainforests as high incoming solar radiation and water availability are the likely drivers of biodiversity (Nee, 2002). Complexity of realization in nature of such stringent conditions of competitive exclusion in according to Def. 1 eliminates the contradictions between the principle and the observed species richness. This is a solution of the biodiversity paradox. The solution of the paradox became possible owing to creation of the white-box models of resource competition and reformulation of the competitive exclusion principle (Kalmykov and Kalmykov, 2013).

Here trying to simplify, generalize and unify the mechanistic Def. 1 we formulate the following definition of the competitive exclusion principle for any number of resource competitors of any nature:

> If a competitor completely prevents any use of at least one essential resource by all its competitors, and itself retains access to all essential resources and ability to use them, then, all other things being equal, all its competitors will be excluded.
>
> — *(Definition 2)*

In formulating a generalized principle of competitive exclusion we relied on the investigated models (Fig. 5). Figure 5a,e represents the case of competitive exclusion, which is consistent with Definitions 1 and 2 of the principle. However, the Definition 1 represents only a direct interspecific competition. In the generalized formulation of the competitive exclusion principle we combine both direct and indirect interspecific competition. Let’s consider a simplest mechanism of indirect interspecific competition (Fig. 5c,g). This is the case when the less fit competitor 2 paradoxically wins the more fit competitor 1. The individual of the species 1 is blocked on all sides and cannot propagate. This is a smallest ecosystem for realization of competitive exclusion in result of *the indirect* conflict of interest. This ecosystem consists of the three microhabitats with two competing individuals. Fig. 5 demonstrates examples of how initial pattern, i.e. initial positioning of individuals on the lattice, size and shape of the lattice, may affect the result.

The necessary condition of the principle is that the competitor “*itself* retains access to all essential resources and ability to use them”. Otherwise, for example, the principle cannot be realized due to extinction of all competitors (Fig. 5d,h) where the both competitors cannot use essential resource for propagation and they both extinct. This is the simplest example of total extinction of the both competitors due to the lack of essential resources.

The generalized form of the competitive exclusion principle allows us to interpret different mechanisms of coexistence in terms of availability of access to essential resource. The both formulations (Def. 1, 2) help to understand a threat to biodiversity that may arise if one competitor will control use of an essential resource of an ecosystem. Today humankind is becoming such a global competitor for all living things (Cincotta et al., 2000; Ehrlich and Ehrlich, 2008; Vitousek et al., 1997).

To expand the theoretical basis for biodiversity conservation we formulate here a principle of competitive coexistence. A theoretical fact of indefinite coexistence of complete competitors on the basis of soliton-like interpenetration of population waves (Kalmykov and Kalmykov, 2013) demonstrates a possibility of formulation of such principle. This ‘soliton-like’ mechanism implements well-timed regeneration of all essential resources and their optimal allocation among competitors. It enables competitors to avoid conflicts of interest. Starting from this mechanism of coexistence of complete competitors, reasoning by contradiction regarding the generalized competitive exclusion principle (Def. 2) and based on the mechanisms of coexistence and extinction of competitors (Fig. 5), we formulate here a second generalized principle – a principle of competitive coexistence, which is also valid for any number of resource competitors of any nature:

> If all competitors retain access to all essential resources and ability to use them, then, all other things being equal and without a global catastrophe, they will coexist indefinitely.
>
> — *(Definition 3)*

Under the global catastrophe we mean a disaster, a fatal disease pandemic, a devastating invasion of predators or herbivores, a mutual destruction. This principle defines the conditions under which a resource competition will not be an obstacle to indefinite coexistence of competitors. Fig. 5b,f represents the case of indefinite competitive coexistence in one closed homogeneous habitat on one and the same limiting resource under constant conditions of environment, without any trade-offs and cooperations. This result confirms the basic hypothesis of this article. Indefinite competitive coexistence implies timely access of all competitors to all essential resources. Such access is the necessary condition for biodiversity conservation and sustainable development.

Previously, by another mechanism, we proved that complete competitors can indefinitely coexist in one closed homogeneous habitat on one and the same limiting resource under constant conditions of environment without any trade-offs and cooperations (Kalmykov and Kalmykov, 2013). Coexistence of the competitors in such conditions is possible due to the mechanism of coexistence which is based on soliton-like behaviour of population waves of competing species. The peculiarities of this mechanism served us as a starting point both for reformulation of the competitive exclusion principle (Def. 1, 2), and for formulation of the principle of competitive coexistence (Def. 3).

## 5. Conclusion remarks

The presented white-box models of ecosystem dynamisc and the formulated here generalized principles of resource competition expand the theoretical basis for biodiversity conservation and sustainable development. In particular, ecological engineering can be considered as designing of sustainable ecosystems that realize optimal use of essential resources.

The methodology of white-box modelling is most effective for the study of complex systems. We consider that the logical deterministic cellular automata is one of the most promising methods of artificial intelligence. This approach allows to provide an automatic deductive inference on the basis of a system of axioms and to get a direct holistic insight into the studied system. Due to the parallel nature of cellular automata this approach is ideal for parallel computing.

## Acknowledgements

This research was partially supported by a reward from the Charity fund of rendering of assistance to scientists “NEW IDEA”.

